# Targeting miRNA for Colorectal Cancer: *In Silico* Identification and Physics-based *De Novo* Modeling of Oncogenic miR-135b for Small Molecule RNA Therapy

**DOI:** 10.1101/2024.06.08.598084

**Authors:** Berat Demir, Lalehan Oktay, Serdar Durdağı

**Affiliations:** Computational Biology and Molecular Simulations Laboratory, Department of Biophysics, School of Medicine, Bahçeşehir University, Istanbul, Türkiye; Department of Molecular Biology and Genetics, Gebze Technical University, Kocaeli, Türkiye; Lab for Innovative Drugs (Lab4IND), Computational Drug Design Center (HITMER), Bahçeşehir University, Istanbul, Türkiye; Molecular Therapy Lab, Department of Pharmaceutical Chemistry, School of Pharmacy, Bahçeşehir University, Istanbul, Türkiye

**Keywords:** miRNA, molecular dynamics simulations, virtual screening, de novo modeling, miR-135, RNA-based therapy

## Abstract

Colorectal cancer is one of the leading causes of cancer deaths worldwide. In the current study, we have identified miR-135b, a microRNA to be differentially expressed in colorectal cancer, through an analysis of the differentially expressed miRNAs from the Gene Expression Omnibus database. Subsequently, the target genes associated with miR-135b were pinpointed, and pathway and functional enrichment analyses were performed to gain a comprehensive understanding of the underlying biological processes involved. A *de novo* three-dimensional model of its tertiary structure was developed for small-molecule targeting at the Dicer cleavage site. Dicer binds to the terminal loop region of the pre-miRNA and cleaves to generate double stranded miRNA duplex. The miRNA duplex is unwound, one of the strands, guide miRNA, is loaded into the RNA-induced silencing complex (RISC) for miRNA-mRNA target interaction and post-transcriptional gene silencing. Following results from molecular docking simulations initiated with the ChemDiv miRNA-targeted small molecule library (∼20.000 compounds), top-scoring compound’s commercial analogues were then searched within the ZINC library using SwissSimilarity. These analogues were docked to the Dicer cleavage site and their optimized docking scores were obtained. These top-scoring molecules were then subject to all-atom molecular dynamics simulations and post-simulation analyses were conducted to assess the dynamic interactions between the miRNA and the selected hit ligands.

## Introduction

Colorectal cancer (CRC) is the third most frequent cancer and the second major cause of cancer deaths in both men and women worldwide.^1^ Even with the advancements in CRC research, 2.5 million new cases of CRC are expected to occur in 2035.^2^ MicroRNAs (miRNAs) are small non-coding RNA molecules which are about 22 nucleotides long and play important regulatory roles in a variety of cellular processes. MicroRNAs carry out their functions by suppressing gene expression at the post-transcriptional level. Biogenesis of miRNAs involves a multi-step process: first, transcription of miRNA genes by RNA polymerase II or III produces primary miRNA (pri-miRNA) transcripts. The Drosha enzyme cleaves the pri-miRNA into hairpin-shaped precursor miRNA (pre-miRNA) in the nucleus, which is subsequently exported to the cytoplasm by Exportin-5. In the cytoplasm, Dicer cleavages the pre-miRNA into a mature miRNA duplex. One strand of the miRNA duplex is then loaded into the RNA-induced silencing complex (RISC), which guides the RISC to the target mRNA through sequence complementarity. The miRNA-RISC complex can cause mRNA degradation or translational repression, which can result in gene silencing. (Figure 1) MicroRNAs have been linked to a variety of biological processes, including development, cell differentiation, proliferation, apoptosis, and stress or pathogen response. The first recognition of the connection between cancer and miRNA dysregulation came from a study by Calin et al.,^3^ identifying that the downregulation of frequently deleted 13q14 region coding two miRNAs was present in the majority of patients with B-cell chronic lymphocytic leukemia.

**Figure 1.**
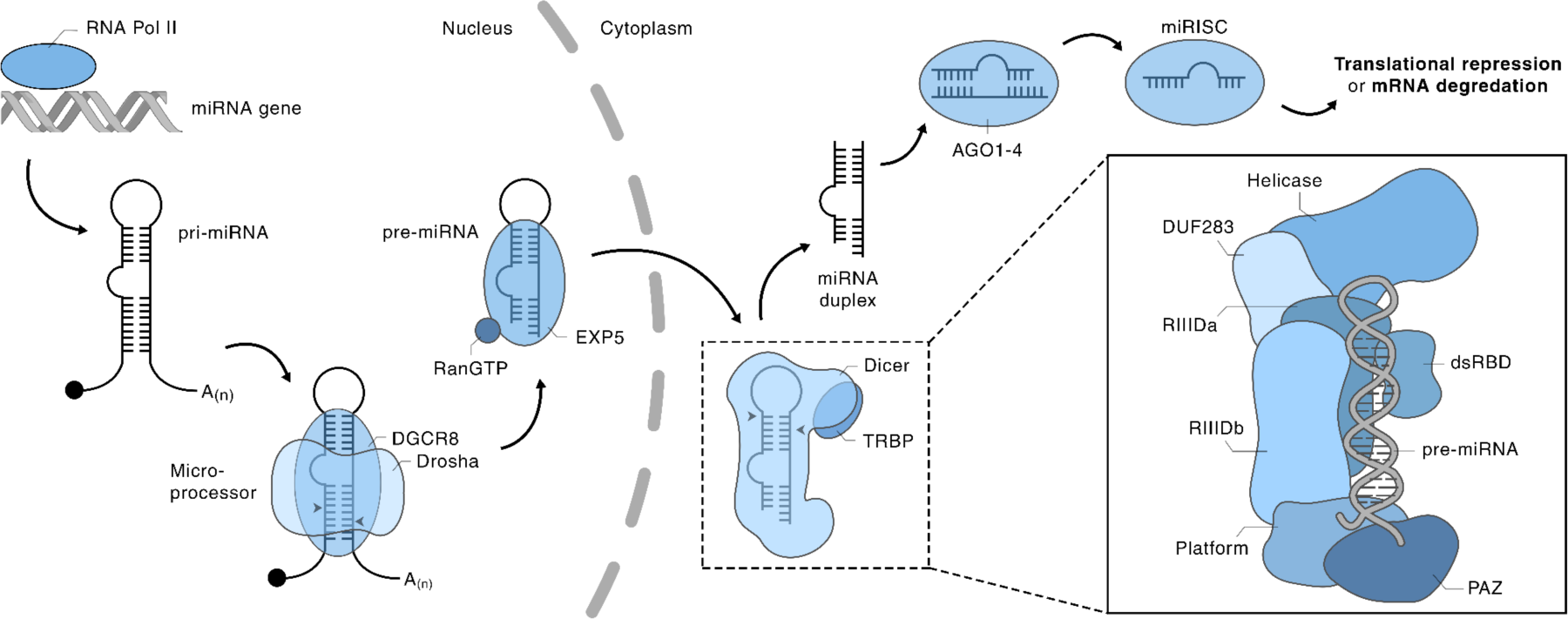
Schematic representation of miRNA biogenesis.

Changes in miRNA level effect a variety of human disorders including cancer, where these alterations become responsible for the initiation and progression of human carcinomas.^4^ The underlying molecular mechanism stems from non-significant modification of gene expression levels which have far-reaching consequences on many genes expressing oncogenic proteins.^5^ Due this ability to regulate multiple genes involved in critical cellular processes, miRNAs have emerged as key players in CRC development and progression.^6^

The uncovering of miRNA expression profiling in human malignant tumors has aided in the unveiling of signature motifs related to diagnosis, prognosis, and treatment response. Furthermore, miRNA profiling is also utilized to identify the downstream oncogenic target proteins and the pathways leading to them.^4^ Specific miRNA expression levels have been linked to CRC initiation, metastasis, and resistance of treatment.^6^ Previously, miR-20a, miR-21, miR-106a, miR-181b, and miR-203 have been identified as having poor prognosis in CRC.^7^ In contrast, some miRNAs, such as miR-143 and miR-145, are downregulated in CRC and play critical roles in inhibiting cell proliferation, inducing apoptosis, and reducing metastasis.^5,6^ Understanding the complex network of miRNAs involved in CRC pathogenesis holds great promise for developing novel diagnostic and therapeutic strategies for CRC treatment.

Targeting the cleavage of the pre-miRNA hairpin sequences is an encouraging therapeutic approach, by selectively inhibiting disease specific miRNA, before they are in their active form. One therapeutic approach, antimiRs, are based on first-generation antisense oligonucleotides (ASOs), are specifically engineered to bind to the mature miRNA and create an inhibition of function.^7^ Further chemical modifications, (i.e. methylation) to antimiRs have shown increase in target specificity, but do have limitations, a major one being the loss of mRNA silencing efficiency.^7^ Furthermore, carrier systems are needed for assistance in their delivery to the site of the tumor. Also, non-small-molecule oligonucleotides are not ideal in the sense of their pharmacodynamic and pharmacokinetic properties.^4^

One great advantage of small-molecule based therapy stems from the fact that they can easily pass through the cellular membranes, drastically increasing the efficiency of the therapeutic efficacy.^8^ This versatility and range of action make small molecules an appealing tool for miRNA-based cancer therapy and an alternative to other invasive therapy approaches.

The current study aims to identify small-molecule inhibitors against oncogenic, differentially expressed miRNAs (DEMs) in CRC. Four DEM datasets, retrieved from the Gene Expression Omnibus (GEO) database, were analyzed to determine the up-regulated DEMs in CRC. Target genes of up-regulated DEMs were identified and pathway and functional enrichment was performed to provide a detailed understanding of the underlying biological processes. After the identification of the up-regulated oncogenic pre-miRNA (miR-135b) in CRC, its tertiary structure was modeled, and docking of the miRNA-targeted small molecule library (∼20.000 compounds) (https://www.chemdiv.com/catalog/structure/mirna-library/) was performed to identify small molecule inhibitors against the pre-miR-135b. Finally, molecular dynamics (MD) simulations and post-MD analysis were performed to qualify the dynamic miRNA-ligand interactions. (Figure 2)

**Figure 2.**
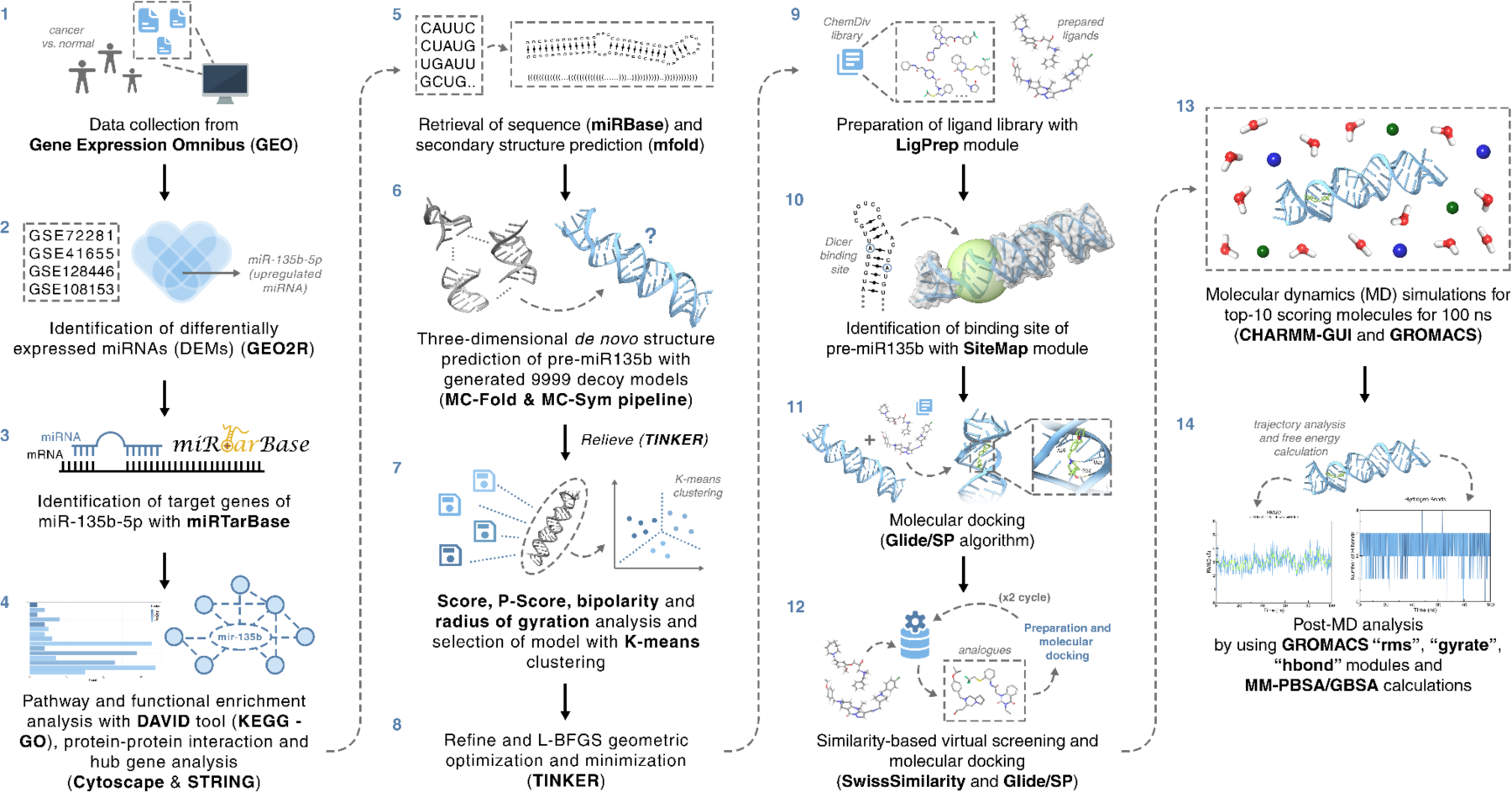
Workflow of the current study. This study integrates identification of DEMs, pathway and functional enrichment analysis, secondary structure and 3D structure predictions, and small molecule screening.

## Materials and Methods

### Data collection

Four expression profiles of miRNAs (GSE72281, GSE41655, GSE128446, GSE108153) were selected from the Gene Expression Omnibus (GEO) database. For a discriminative search of the datasets, "miRNA” and “colorectal cancer” were used as keywords. GSE72281, using GPL18058 platform (Exiqon miRCURY LNA microRNA array, 7^th^ generation), contained 6 colorectal cancer samples and 6 normal samples; GSE41655, using GPL11487 platform (Agilent-021827 Human miRNA Microarray), contained 33 colorectal cancer samples and 15 normal samples; GSE128446, using GPL14767 platform (Agilent-021827 Human miRNA Microarray G4470C), contained 18 colorectal cancer samples and 4 normal samples and GSE108153, using GPL19730 platform (Agilent-046064 Unrestricted Human miRNA V19.0 Microarray), contained 21 colorectal cancer samples and 21 normal samples.

### Identification of up-regulated differentially expressed miRNAs

Colorectal cancer and control groups were specified for each dataset. For the comparison of the samples, R-based web tool GEO2R (using “limma” package) was used.^9,10^ DEMs were screened for each dataset according to the cut-off value of the adjusted p-value < 0.05 [using Benjamini & Hochberg (False discovery rate)].^11^ For more specific and up-regulated miRNA results, the cut-off value of log2FC > 2 was set. After analyzing all datasets, overlapping miRNAs for all datasets were identified by a Venn diagram.

### Identification of target genes of DEMs

Target prediction of DEMs was carried out by miRTarBase^12^, an experimentally verified miRNA-target interactions database. mRNA genes identified as targets were recorded.

### Pathway and functional enrichment analysis

The genes identified as the targets of DEMs were submitted to the Database for Annotation, Visualization and Integrated Discovery (DAVID)^13^ online tool for Kyoto Encyclopedia of Genes and Genomes (KEGG)^14^ pathway enrichment and Gene Ontology (GO) functional enrichment analysis^15^ with EASE score (p-value) < 0.05 as the cut-off. DAVID proposes a complete set of functional annotation tools to solve biological significance of lists of genes. Gene Ontology (GO) offers functional enrichment from 3 different aspects for target genes: Biological process, cellular component, and molecular function.

### Protein-Protein Interaction (PPI) construction and hub gene analysis

The PPI network of target genes was constructed using the Search Tool for the Retrieval of Interacting Genes (STRING)^16^ with an interaction score of 0.4 (medium confidence). STRING is a database of known and predicted PPI, including direct (physical) and indirect (functional) relationships. The PPI data retrieved from the STRING database were visualized and analyzed in Cytoscape^17^, which is an open-source tool for integration, visualization, and analysis of biological networks. Top-10 important hub genes were identified by cytoHubba^18^, a Cytoscape plugin, using the maximal clique centrality (MCC) algorithm^18^.

### miRNA 3D structure prediction

The pre-miRNA stem-loop sequence of the differentially expressed and up-regulated miRNA was obtained from the miRBase^19^ database. The secondary structure of the miRNA was identified by the physics-based tool mfold^20^, which works by predicting the minimum free energy. Tertiary structure of the miRNA was predicted using the MC-Fold / MC-Sym^21^ web-hosted server pipeline. First, the secondary structure of the miRNA obtained with mfold was masked using the MC-Fold tool in dot-bracket (Vienna) format. The tertiary structure of the miRNA was then predicted by the MC-Sym server. The MC-Sym web server, uses a fragment-based approach by the Nucleotide Cyclic Motifs (NCM) fusion process to generate 3D decoy models. To remove steric clashes and adjust the positions of the backbone atoms, generated decoy models were relieved using the steepest descent optimization algorithm with the TINKER Molecular Modeling Package. Parameters like *“score”*, “*P-score”*, *“bipolarity”* and *“radius of gyration”* values were calculated to compare all decoy models. The *“score”* parameter computes the internal energy of the decoy models. The *“p-score”* parameter measures the A-RNA likeness of decoy models and their quality by using the positions of the phosphate atoms.^21^ The bipolarity parameter relates the coplanarity of the RNA base-normal vector arrangements with the validation of RNA secondary structure.^22^ Here, the radius of gyration indicates the size of the folded RNA structures, by using the coordinates of the structures of RNA.^21^ Cut-off values for the parameters were determined as *“score”* < -50, *“p-score”* < -30, “*bipolarity”* > 0.8 and 18.3 < *“radius of gyration”* < 26.4. Models determined suitable were selected for k-means clustering with 5 clusters calculated using the default framework. Among the five clusters, the models with the lowest scores were determined, and the model with the lowest score of the cluster with the most models was selected. Refinement and L-BFGS quasi-newton optimization were applied to adjust the energy and geometric optimization of the selected decoy model by using TINKER Molecular Modelling Package.^23^

### Ligand preparation

The miRNA targeted small molecule drug library containing ∼20.000 small molecules was retrieved from the ChemDiv database which was curated by (i) using ML-based models from ECFP fingerprints derived from previously identified miRNA functional modulators to screen the ChEMBL29 database, (ii) applying the PAINS filters, (iii) structural diversity picking (https://www.chemdiv.com/catalog/structure/mirna-library/). The downloaded library was prepared with the OPLS3e force field^24^ using the LigPrep module of the Maestro molecular modeling package (Schrödinger Release 2018-1, LigPrep, Schrödinger). The pH value was set to 7.4 ± 0.5 for the assignment of suitable protonation states.

### Selection of the binding site

Ligand binding site of the miRNA was predicted with the SiteMap module^25^ of the Maestro molecular modeling package. The selected site was further validated by aligning the mature miRNA sequence to that of the pre-miRNA and comparing the omitted residues to yield the pre-miRNA.

### Molecular Docking

Prepared miRNA targeted small molecule library was docked to the predicted 3D structure of the pre-miRNA. The Glide/SP (standard precision)^26^ algorithm implemented in the Maestro molecular modeling package using flexible ligand sampling. The receptor grid box was derived from the Dicer binding site Cartesian coordinates provided by the SiteMap^27^ module.

### Similarity Search with Ligand-Based Virtual Screening (LBVS)

Top-5 ranking molecules with best docking scores were selected and the similarity search for each molecule was conducted using the SwissSimilarity server (http://www.swisssimilarity.ch/), which is a web-based tool used that performs similarity-based screening from input ligand on large chemical libraries. The class of molecules to be searched from was set as “commercial” and the ZINC drug-like compounds library was selected. To calculate the combined score between selected molecules and compounds in the library, FP2 fingerprints and ElectroShape-5D (ES5D) vector combined screening methods were used. The combined score indicates calculation from similarity values obtained from FP2 and ES5D methods and it is significantly better for drug-like compounds than calculation with these methods separately. After the searching process, all structural analogues found were docked to the previously generated 3D pre-miRNA employing the Glide/SP docking algorithm. This process was further repeated for two cycles for resulting molecules which had docking scores better than their parent analogue molecule.

### Molecular Dynamics (MD) simulations

MD simulations were performed by GROMACS-2022.^28^ The pre-miRNA - ligand complex was prepared using the CHARMM-GUI web-based tool^29^ to build and prepare the system as an input for MD tools. RNA-ligand complex topology files were generated using the Amber OL3 RNA force field^30^ for RNA and the General Amber Force Field 2 (GAFF2)^31^ for ligand. The complex was then plunged in a periodic water box with 10 Å edges from the complex, composed of TIP3P waters and 0.15M Na^+^, Cl^-^ ion concentration. Energy minimization was performed on the whole system using the steepest descent algorithm for 1000 steps. Hydrogen bond lengths were maintained with respect to the LINCS algorithm.^32^ The system was further equilibrated employing the NPT ensemble with the Nose-Hoover temperature coupling ^33,34^ at 310 K and Parrinello-Rahman isotropic pressure coupling.^35^ Production MD simulations were run for 100 ns with 3 replicates for each of the RNA-ligand complexes. The Verlet cutoff scheme^36^ was employed, where the long-range interactions were predicted with the Particle Mesh Ewalds (PME) scheme^37^ and the short-range interactions were calculated at 9 Å. Post-simulations, RMSD, radius of gyration, potential energy, solvent accessible surface area and H-bond analyses were performed with GROMACS “rms”, “gyrate” and “hbond” modules. All RNA-ligand complexed simulations were repeated 3 times, and simulations with frames exceeding 15 Å RNA fit to ligand RMSD were discarded.

### MM-PBSA/GBSA Binding Free Energy Calculations

Binding free energy was calculated using the *gmx_MMPBSA* tool with molecular mechanics Poisson-Boltzmann and generalized Born surface area (MM-PB(GB)SA) methods. GB-Neck2 (igb=8) was set as the solvation model for GB calculations, and PB model was set as default (ipb=2). Internal (solute) dielectric constant was set to 1 and external (solvent) dielectric constant was set to 78.5. All computations were carried out with a NaCl concentration of 0.15 M and a solvent probe radius set at 1.4 Å. The interval value was set to 1, calculations were run for all frames within 100 ns simulations. All procedures were applied individually to all remaining pre-miR135b - ligand simulations. Enthalpy scores using both methods were collected.

## Results and Discussion

### miR-135b is differentially expressed in colorectal cancer

The data with colorectal cancer and control groups, GSE72281, GSE41655, GSE128446 and GSE108153, were retrieved from the GEO database. GEO2R, R-based web tool, was used to identify the up-regulated differentially expressed miRNAs (DEMs) in GSE72281, GSE41655, GSE128446 and GSE108153 datasets. For selection of the significantly up-regulated DEMs, the adjusted p-value < 0.05 and log2FC > 2 were used as cut-off criteria. In total, 17 miRNAs in the GSE72281 dataset, 15 miRNAs in the GSE41655 dataset, 126 in the GSE128446 dataset, and 3 miRNAs in the GSE108153 dataset were identified as up-regulated DEMs. All identified miRNAs were statistically significant due to adjusted p-value < 0.05. The volcano plot of miRNAs in GSE72281(Figure 3A), GSE108153 (Figure 3C), GSE41655 (Figure 3C) and GSE128446 (Figure 3D) representing the expression of miRNAs in colorectal cancer (compared with adjacent tissue, normal tissue and normal mucosa) were plotted. Venn’s diagram analysis was used to obtain the overlapped up-regulated DEMs in all datasets. The analysis showed that one up-regulated miRNA (hsa-miR-135b-5p) was common in all the datasets (Figure 3E). Furthermore, in a recent study evaluating the gene expression of genes BCL2, BAX, CASP3, CASP9, MMP2, MMP9 and 14 related miRNAs in colorectal cancer identified eight of these miRNAs, including miR-135, to be upregulated.^38^

**Figure 3.**
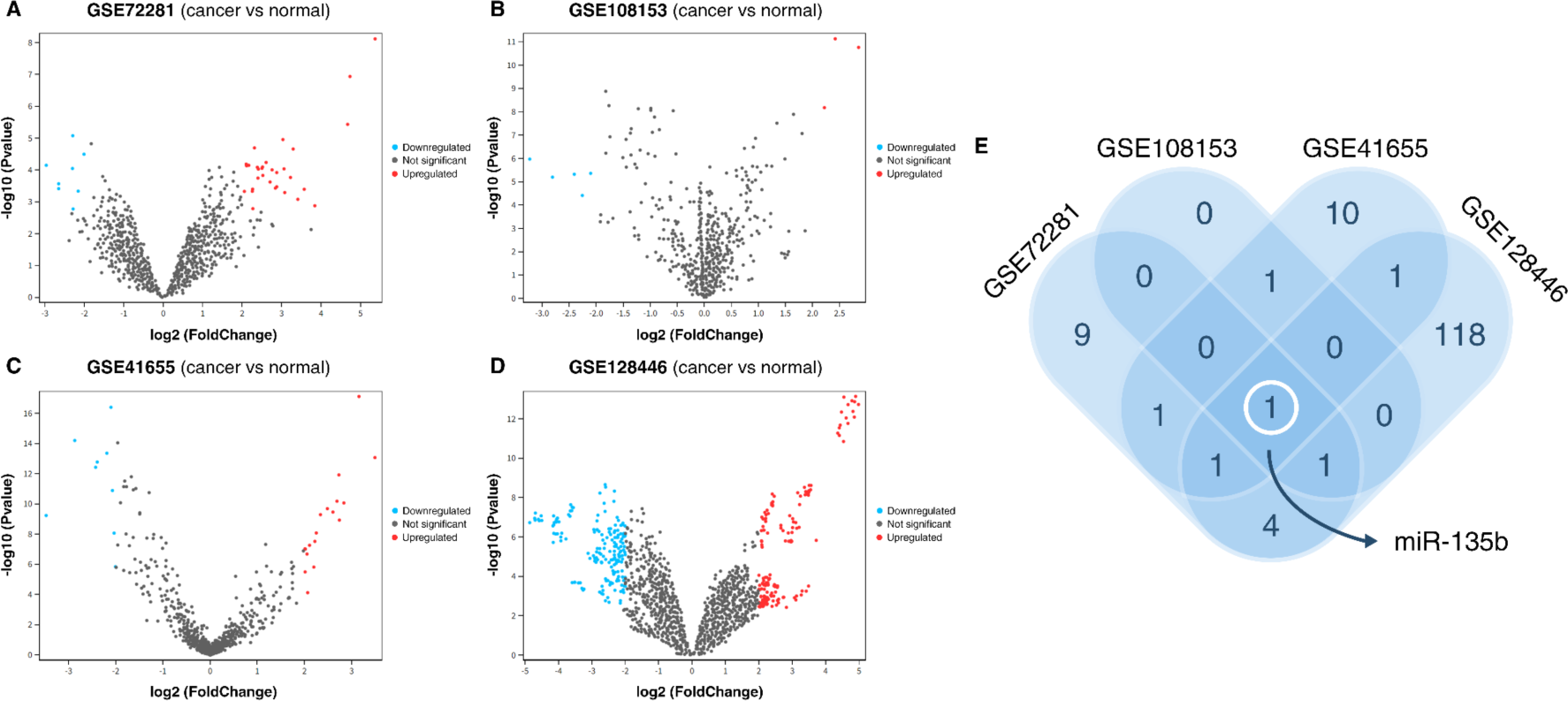
Identification of differentially expressed miR-135b in colorectal cancer. Volcano plots of miRNAs in **(A)** GSE72281, **(B)** GSE108153, **(C)** GSE41655 and **(D)** GSE128446 representing the expression of miRNAs in colorectal cancer (compared with adjacent tissue, normal tissue and normal mucosa). **(E)** Venn’s diagram analysis of all up-regulated DEMs overlapped.

### GO functional and KEGG pathway analysis of target genes of miR-135b

GO functional and KEGG pathway analysis were done in the DAVID database. Retrieved miR-135b-5p target genes were submitted to the DAVID database. For strong enrichment, the EASE score (p-value) threshold was selected as <0.05. Most miR-135b target genes were enriched in “regulation of transcription from RNA polymerase II promoter”, “negative regulation of transcription from RNA polymerase II promoter”, “ positive regulation of transcription from RNA polymerase II promoter” as biological processes (GO_BP) (Figure 4A); “protein binding”, “RNA polymerase II core promoter proximal region sequence-specific DNA binding”, “RNA polymerase II transcription factor activity, sequence-specific DNA binding” as molecular functions (GO_MF) (Figure 4B); and “nucleus”, “cytosol”, “cytoplasm” as cellular components (GO_CC) (Figure 4C). The results showed that the most significant KEGG pathways for miR-135b-5p target genes were “signaling pathways regulating pluripotency of stem cells”, “TGF-beta signaling pathway”, “Colorectal cancer” and “Pathways in cancer”. (Figure 4D)

**Figure 4.**
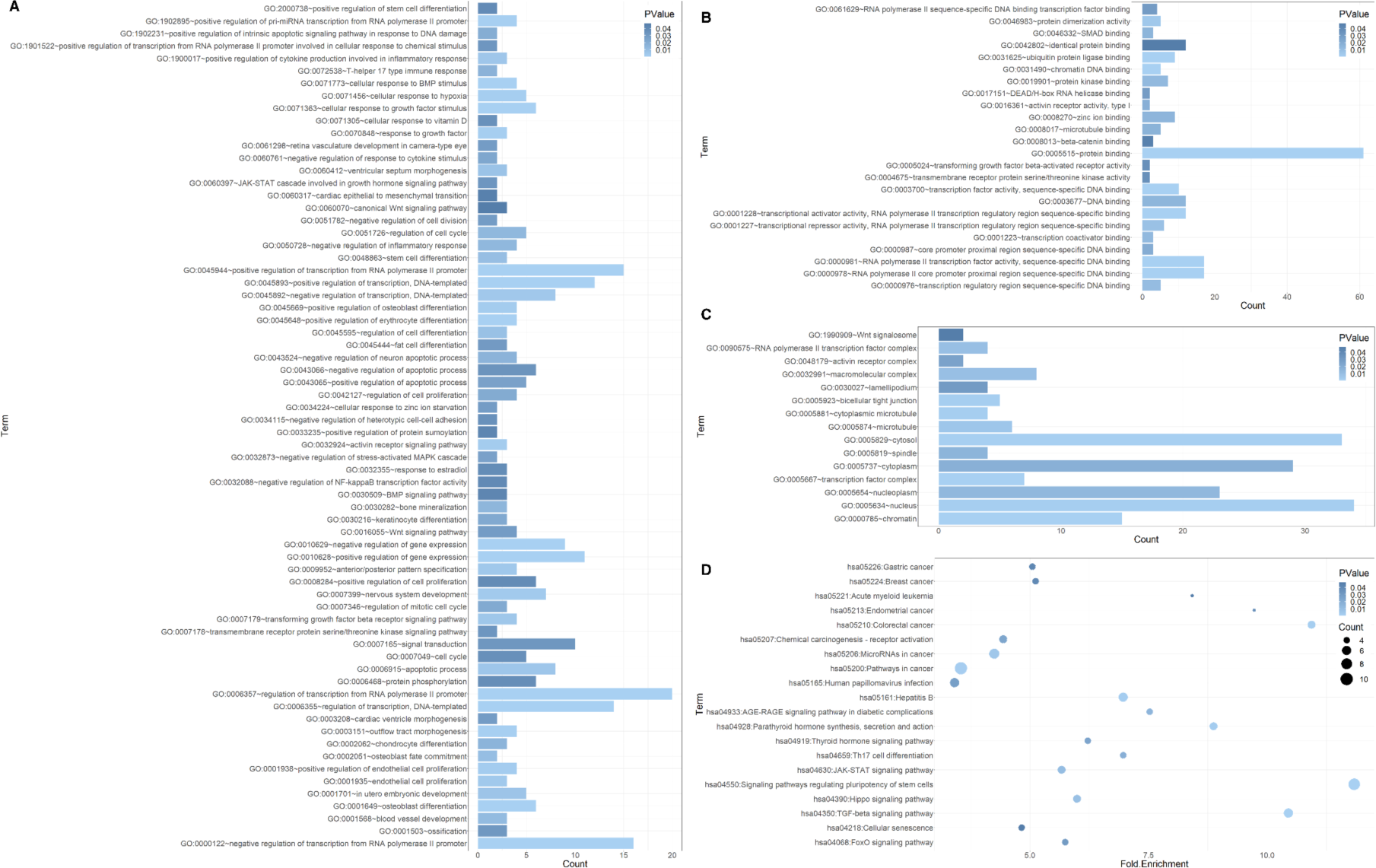
GO functional and KEGG pathway analysis of target genes of miR-135b. **(A)** GO_BP (Biological Process), **(B)** GO_MF (Molecular Function) and **(C)** GO_CC (Cellular Components) functional enrichment plots. **(D)** KEGG pathway enrichment plot.

### PPI analysis reveal STAT3, MYC, HIF1A, FOXO1, KLF4, RUNX2, KRAS, TGFBR1, MEF2C, and STAT6 hub genes associated with miR-135b-5p

The PPI network of miR-135b-5p targets constructed using the STRING database was implemented in Cytoscape^17^. cytoHubba^18^ was used to identify hub genes of miR-135b and calculate the nodes’ scores (Figure 5A). Using a maximal clique centrality (MCC) topological analysis algorithm, top-10 important hub genes were identified. These genes were ranked as STAT3, MYC, HIF1A, FOXO1, KLF4, RUNX2, KRAS, TGFBR1, MEF2C, and STAT6 from highest score to least, respectively. (Figure 5B) In addition, among these 10 hub genes, MYC, KRAS and TGFBR1 genes were enriched in "colorectal cancer" according to KEGG pathway analysis as well. In a previous study conducted by Li et al.,^39^ miR-135b was recognized as a facilitator of tumor growth in CRC cells. This occurred by enhancing their proliferation and inhibiting apoptosis through the reduction of TGFBR2 expression. However, the study did not provide any information on the expression levels of TGFBR1.^39^ Another study by Nagel et al., implicates miR-135b in colorectal cancer progression through downregulation of the tumor suppressor APC gene.^40^

**Figure 5.**
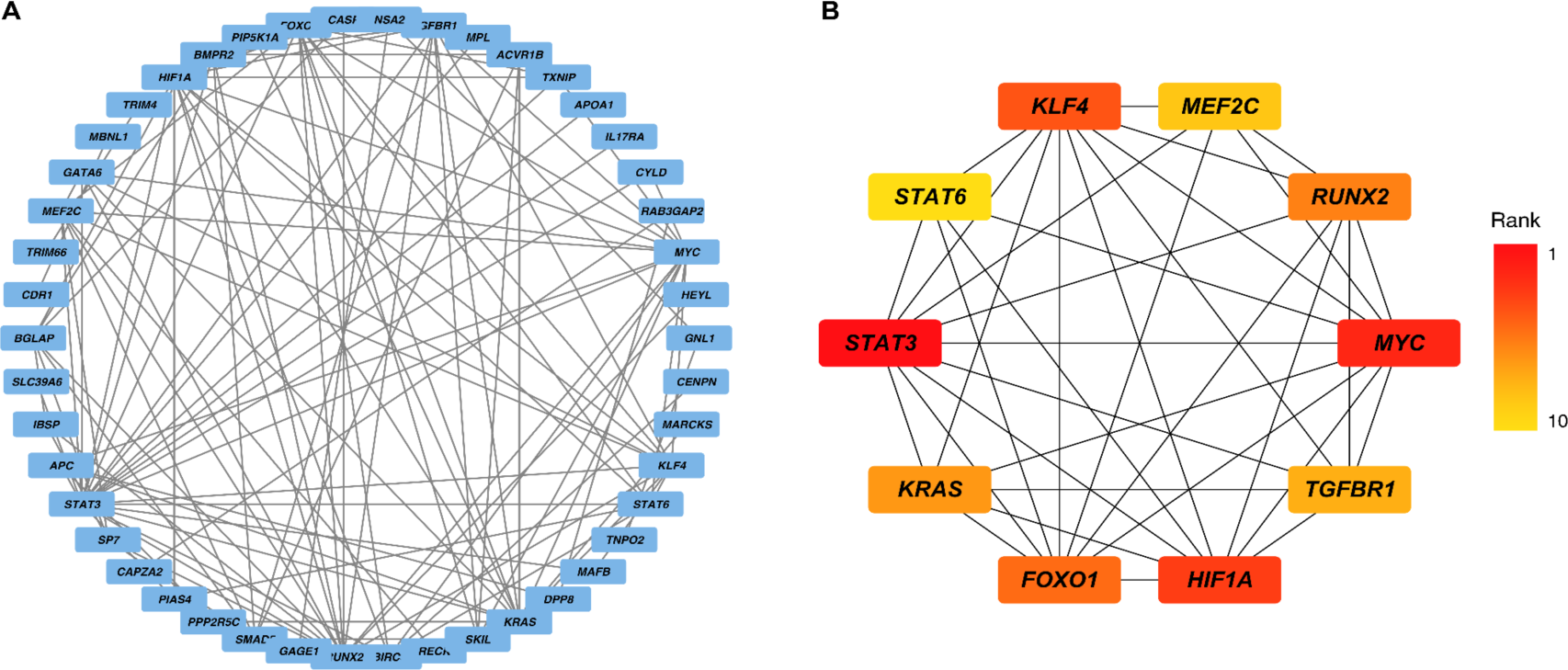
miR-135b-5p PPI analysis and prediction of hub genes. **(A)** PPI network of miR-135b-5p from STRING. **(B)** Top 10 important hub genes ranked as STAT3, MYC, HIF1A, FOXO1, KLF4, RUNX2, KRAS, TGFBR1, MEF2C, and STAT6 from highest score to least, respectively.

### *De novo* modeling of pre-miR-135b Tertiary Structure Uncovers the 3D Dicer Binding Domain

The tertiary structure of pre-miR135b, given in Figure 6A, was predicted using mfold and subjected to the MC-Fold/MC-Sym pipeline. For the convenience of de novo modeling, only the region between nucleotides 13 and 67 (63 base pairs) including the hairpin loop region, was used for prediction of the 3D structure (Figure 6A). 9999 decoy models were then generated for 3D structure prediction with MC-Sym. Decoy models were relieved using the TINKER Molecular Modeling Package^23^ and “score”, “p-score”, “bipolarity” and “radius of gyration” values of the structures were calculated and models fitting within the cut-off intervals of these parameters were selected. The generation of RNA structural ensembles, particularly if there is no experimental validation, has its challenges. Using molecular mechanics-based forcefields falls short on judging the correctness of a representative model, since the force-fields can have limited outlook of the overall perspective and fail to assess imperfections in models refined with MC-Sym. Thus, parameters with customizable intervals represent the optimal way for selecting the most correct model from the ensemble set. Chosen models were clustered with the k-means clustering algorithm into 5 clusters. Clusters 1 to 5 were populated with 10, 8, 7, 6 and 5, respectively. Each cluster representative along with its model number, p-score, score, radius of gyration and bipolarity is presented in Table 1. The 1713^th^ model, denoted model 1713, which has obtained the lowest score value in the cluster with the most models (cluster 1), was selected. Model 1713 was refined and L-BFGS Quasi-Newton geometric optimization was performed by using TINKER Molecular Modeling Package.^23^ The 3D pre-miR-135b and the nucleotides included in the mature miRNA are shown in Figure 6. The Dicer-binding site, identified from the 3D structure of the modeled pre-miR-135b (Figure 6B) encompasses bases U23, G24, A25, U26, A37, A38, U40, C41 and A42. This region was preferred amongst the generated SiteMap since it corresponds to the hairpin 2nt-3’ overhang region recognized by Dicer.

**Figure 6.**
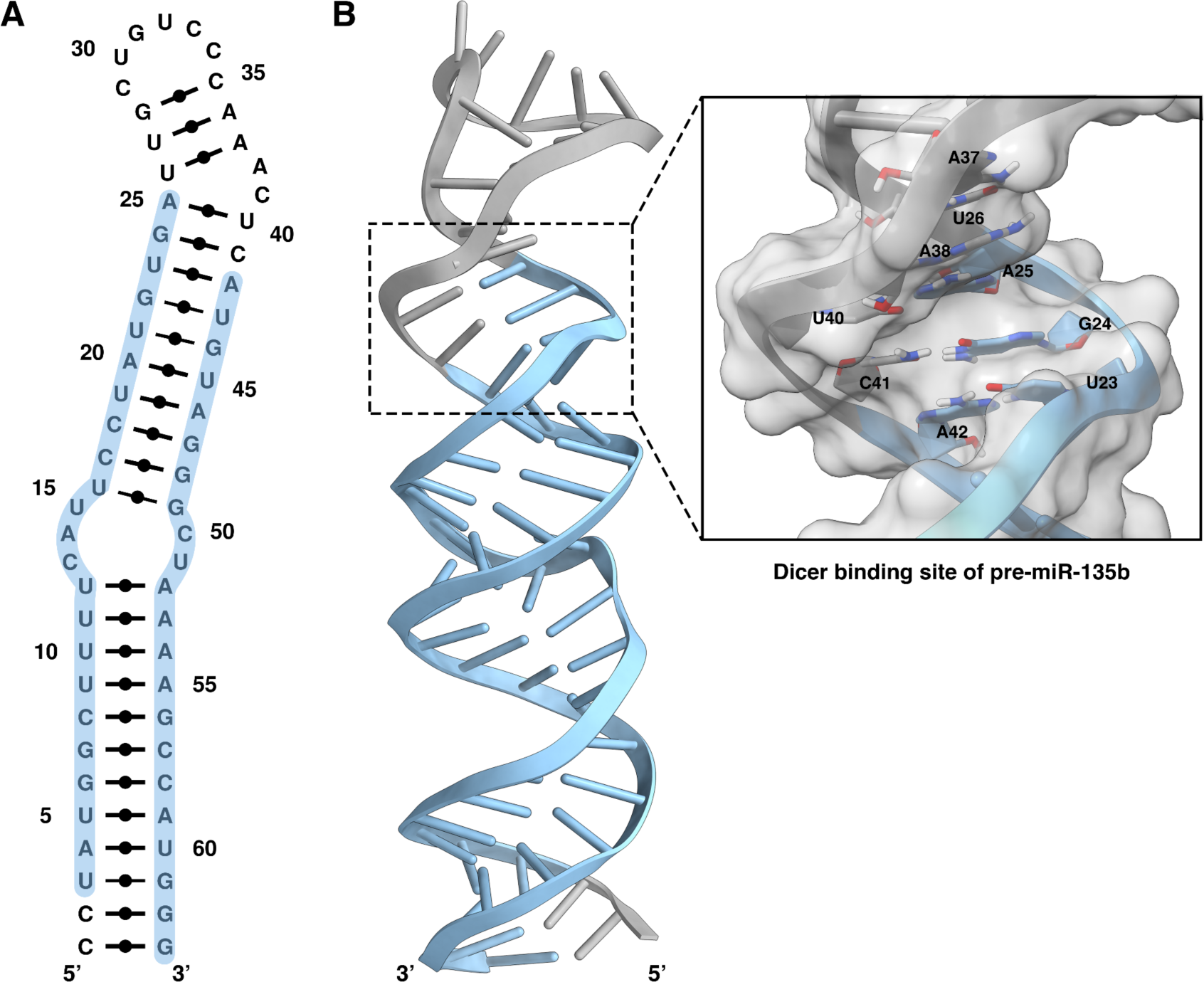
Structure of pre-miR135b and identification of the Dicer binding site. **(A)** Secondary structure of selected shortened sequence of pre-miR135b, nucleotides highlighted in blue indicate mature miRNAs, **(B)** 3-D representation of selected pre-miR135b model and Dicer binding site of pre-miR135b.

**Table 1.**
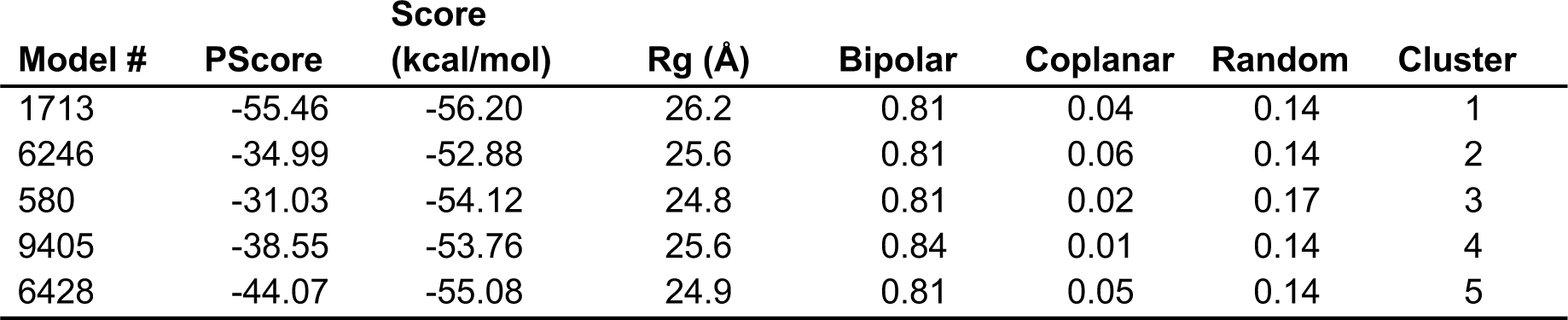
PScore, score, radius of gyration and bipolarity analysis of the top-scored models in clusters.

| Model # | PScore | Score<br>(kcal/mol) | Rg (Å) | Bipolar | Coplanar | Random | Cluster |
| --- | --- | --- | --- | --- | --- | --- | --- |
| 1713 | -55.46 | -56.20 | 26.2 | 0.81 | 0.04 | 0.14 | 1 |
| 6246 | -34.99 | -52.88 | 25.6 | 0.81 | 0.06 | 0.14 | 2 |
| 580 | -31.03 | -54.12 | 24.8 | 0.81 | 0.02 | 0.17 | 3 |
| 9405 | -38.55 | -53.76 | 25.6 | 0.84 | 0.01 | 0.14 | 4 |
| 6428 | -44.07 | -55.08 | 24.9 | 0.81 | 0.05 | 0.14 | 5 |

### Molecular Docking and Similarity-Based Virtual Screening Identifies Potential Hit Molecule Candidates for pre-miR-135b Maturation Inhibition

The quest for small-molecule inhibitors targeting precursor miRNAs mainly focuses on specific secondary structures formed by single-stranded regions like loops and bulges, in addition to double-stranded areas, creating distinct RNA binding pockets.^41^ Nonetheless, small molecule-based inhibition strategies of the pre-miRNA maturation process is a well-established approach.^42–45^. According to the miRBase database, the mature miR-135b-5p sequence spans from nucleotides 16 to 38, while the mature miR-135b-3p sequence extends from nucleotides 55 to 76. Consequently, it can be deduced that both the 38^th^ and 55^th^ nucleotide residues are located within the region where Dicer binds. Hence, this specific region has been identified as the Dicer binding region. In this context, the miRNA targeted small molecule library obtained from the ChemDiv database (https://www.chemdiv.com/catalog/structure/mirna-library/) was used for docking-based virtual screening for potential pre-miR-135b/Dicer complex formation inhibitors. Docking scores were recorded and the ligands with the lowest-energy compounds (i.e., high docking scored compounds) were subjected to similarity-based screening. The SwissSimilarity server (http://www.swisssimilarity.ch/) is used to determine the analog structures of the 5 molecules with the high docking scores. As a result of the screening with the “commercial” ZINC drug-like compounds library, analogs of 5 high-scoring molecules were obtained and docking was carried out again with the Glide/SP docking algorithm for the obtained 1663 analog molecules. Then, the molecules that passed the docking scores of the top-5 molecules were recorded and similarity search was carried out again in the SwissSimilarity web-tool for these molecules. Resulting 800 analogues were docked to the pre-miR-135, as well. All docking results for ChemDiv molecules and molecules obtained from the SwissSimilarity workflow were ranked based on their docking scores and the top-10 molecules were recorded. Four of these molecules were from ChemDiv miRNA targeted small molecule library and the remaining 6 molecules were from the ZINC drug-like library obtained from the SwissSimilarity search effort. (Table 2) The resulting 6 molecules from the SwissSimilarity search of the ZINC drug-like library have obtained better docking scores than their initial analogues (Table 2), proving that screening workflows can be extended and ameliorated with the addition of a similarity-based workflow.

**Table 2.** Docking score of selected molecules from Glide/SP.

| Compound ID | Docking<br>Score<br>(kcal/mol) | Library | Cycle<br>count | Derived from<br>Compound ID | Derived from<br>Compound<br>Library |
| --- | --- | --- | --- | --- | --- |
| G700-0411 | -9.93 | ChemDiv | 0 |  |  |
| P101-1799 | -9.93 | ChemDiv | 0 |  |  |
| S986-0170 | -9.93 | ChemDiv | 0 |  |  |
| ZINC000096491549 | -9.81 | ZINC | 2 | CM2432-0321 | ChemDiv |
| C336-0173 | -9.77 | ChemDiv | 0 |  |  |
| ZINC000097784705 | -9.70 | ZINC | 2 | CM2432-0321 | ChemDiv |
| ZINC000015062379 | -9.59 | ZINC | 2 | CM2432-0321 | ChemDiv |
| ZINC000014882460 | -9.59 | ZINC | 2 | CM2432-0321 | ChemDiv |
| ZINC000091746401 | -9.56 | ZINC | 2 | CM2432-0321 | ChemDiv |
| ZINC000091965114 | -9.55 | ZINC | 1 | CM2432-0321 | ChemDiv |

### MD Simulations show stability in RNA Shape Upon Ligand Binding

To determine the structural changes induced pre-miR-135b upon ligand binding, 3 replicates of MD simulations (100 ns) were conducted for the top-10 scoring molecules resulting from screening the ChemDiv library and further SwissSimilarity cycles. Two of the selected ten compounds deviated and flew away from the designated binding pocket in all three replicate MD simulations and were thereof not considered in the previously mentioned post-MD calculations. Also, in some of the surviving compound replicate simulations, over 15 Å deviations were detected and thus these simulations were also discarded. A representative frame drawn from the ZINC97784705-RNA simulation demonstrates how the docked ligand should reside in the postulated dicer binding site. (Figure 7A) 2D ligand interaction diagram in Figure 7B, highlights the bonded interactions conserved during most of the simulation time. Post-simulation normalized and averaged over replicas RNA fit to ligand RMSD calculations (Figure 7C, right) have revealed the probability of the RMSD fluctuation during 100 ns of simulations. Here, ZINC library compound ZINC97784705 from the SwissSimilarity workflow shows the lowest RNA-ligand RMSD overall for a period of 100 ns and 1000 frames. Compound G700-0411 has the largest RMSD fluctuation, owing to the slight translocation of the ligand in the binding pocket. RNA-ligand RMSD of ZINC97784705 (Figure 7C, left) was also plotted to assess time dependent RMSD progression. Although RNA has a highly volatile nature in MD simulations^46^, we can observe the relative stability of the ZINC97784705-RNA complex.

**Figure 7.**
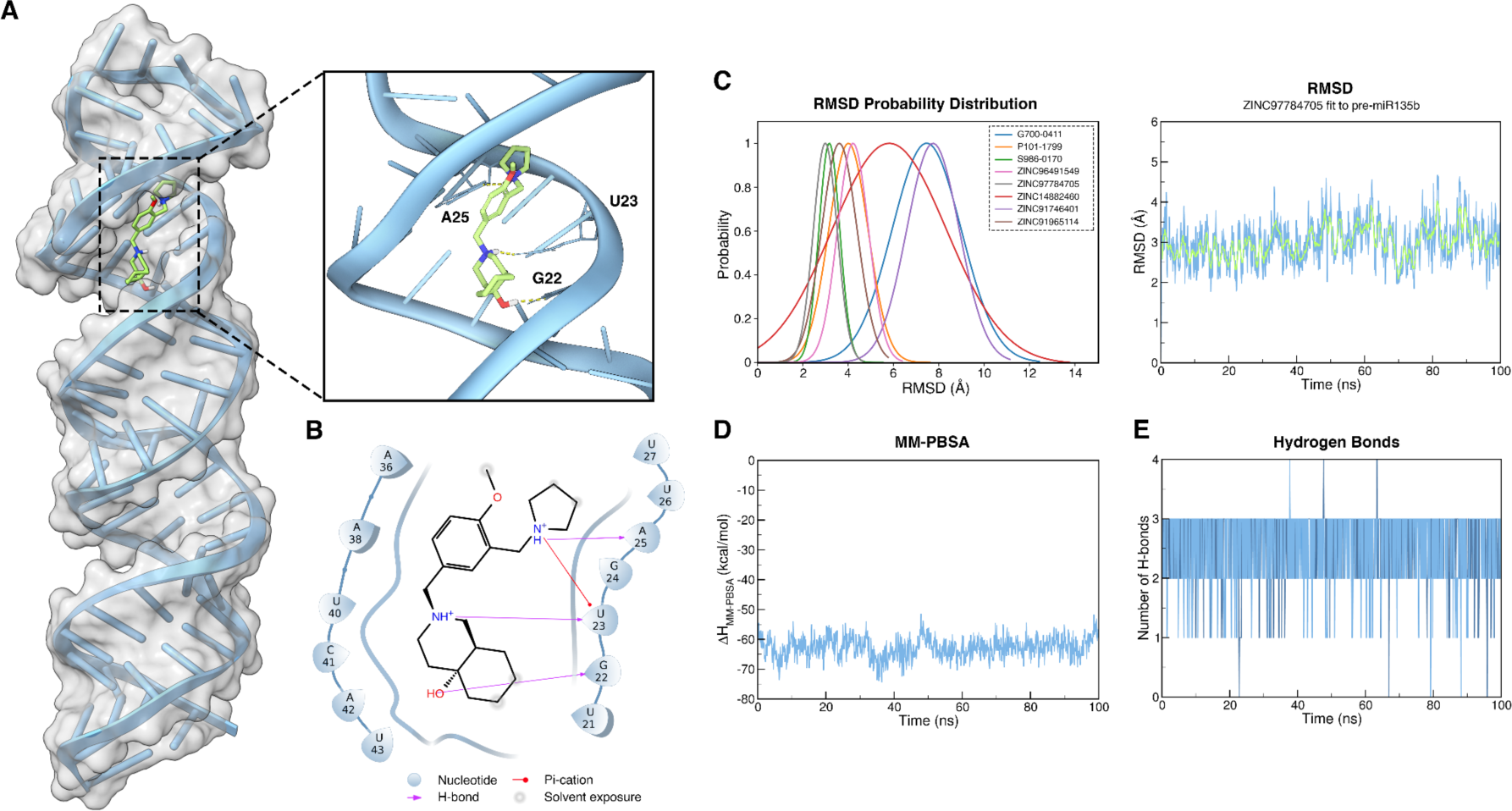
Post-MD Analysis for ZINC97784705. **(A)** 3D representation of pre-miR135b and ZINC97784705 complex, **(B)** 2-D interaction diagram for ZINC97784705, **(C)** ZINC97784705 fit pre-miR135b RMSD probability distribution and RNA-ligand RMSD plot **(D)** MM-PBSA scores per frame and **(E)** Time dependent hydrogen bond analysis.

ΔH_MM-PBSA_ scores per frame for the ZINC97784705-RNA complex (Figure 7D) were further plotted to validate that the RMSD stability of the RNA-ligand complex was indeed related to energetically favorable interactions. It can be observed that the ΔH_MM-PBSA_ stays between -60 and -70 kcal/mol. Time-dependent hydrogen bonding analysis (Figure 7E) quantifies the previous observation, demonstrating the 2 to 3 hydrogen bonds formed between ligand and protein, observable in all replicas, namely between bases G22, U23 and A25 as shown in 3D in Figure 7A. ΔH_MM-GBSA_ for all hit compound-RNA complexes were calculated for all trajectory frames (Table S1) and for the last 501 frames of simulations. (Table S2)

Assessing the dimensions and flexibility of 3D RNA structures are crucial for conceptualizing the arrangement of secondary structures and their interactions with small molecules or proteins. As per the Flory theory^47^, the radius of gyration for RNA increases proportionally to its number of nucleotides such that;

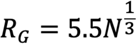

where symbolizes the number of nucleotides and the coefficients used were recommended in the MC-Sym workflow. In the case studied above, our proposed 3D miR-135b model has 63 nucleotides, therefore an expected value of ∼22 Å. The average of all miRNA-ligand complexes fluctuates around 26 Å (Table 3), which correlates with the (26.2 Å) of the initial starting structure. From ligand-bound simulations, we have assessed that ligand binding does not necessarily influence the size and compactness of the miRNA. (Table 3)

**Table 3.** Radius of gyration analysis of ligand-bounded pre-miR135b structures.

| Compound ID | Minimum Rg (Å) | Maximum Rg (Å) | Average (Å) |
| --- | --- | --- | --- |
| G700-0411 | 22.26 | 29.76 | 26.13±1.26 |
| P101-1799 | 22.57 | 28.50 | 25.47±0.96 |
| S986-0170 | 23.78 | 29.05 | 26.03±0.91 |
| ZINC000096491549 | 22.95 | 28.75 | 25.55±0.95 |
| C336-0173 | 22.33 | 29.69 | 26.03±1.28 |
| ZINC000097784705 | 23.59 | 28.71 | 25.98±0.87 |
| ZINC000015062379 | 23.55 | 29.41 | 26.29±0.88 |
| ZINC000014882460 | 23.18 | 28.47 | 26.04±0.92 |
| ZINC000091746401 | 23.08 | 30.09 | 26.60±1.12 |
| ZINC000091965114 | 22.50 | 29.28 | 25.78±1.00 |

## Conclusions

Utilizing the advantages of small-molecule inhibitors to target miRNAs is a relatively new strategy for treating human diseases. This approach combines the superiority of small-molecule therapeutics with targeting disease-specific miRNAs, providing tailor-made therapeutic solutions. The targeted mechanism is that of the Dicer-mediated miRNA maturation process. The Dicer double-stranded RNA-binding domain (dsRBD) domain recognizes either the GYM motif or a single-nucleotide bulge near the terminal end of the hairpin-loop.^48^ After Dicer cleavage, most miRNAs will bind to target mRNAs’ 3’ UTR region and regulate its translation.^49^ Therefore, hampering the Dicer-pre-miRNA interaction stands out as an efficient strategy for inhibiting miRNA mediated translational dysregulation. From statistical analysis performed on the DEM datasets from the GEO database, miR-135b stands out as being differentially overexpressed in all 4 datasets (Figure 3E). Additionally, in a recent investigation of miRNA expression profiles in colorectal cancer identifies miR-135 as one of the differentially upregulated miRNAs. Utilizing the tertiary structure of miRNAs for small-molecule inhibition through physics-based approaches have many advantages; namely that the characterization of the pre-miRNA-dicer binding pocket is very well defined, aiding in the correct screening of small-molecules based on their electrostatic fitting. The secondary and tertiary structure identification of miRNAs have pivotal importance in locating the potential binding pocket. Regarding small-molecule RNA targeting, there is still a lack of standardized methodology.^50^ Our study has tested a comprehensive approach to *de novo* modeling, Dicer binding site identification and targeted small-molecule screening with RNA, yielding potential hits targeting miR-135b for colorectal cancer treatment. Specifically, we identify ZINC97784705 as a potential miR-135b-targeted colorectal cancer therapeutic strategy, containing a pyrrolidine group observed in pacritinib and futibatinib, both FDA-approved anticancer drugs.^51^

## Supporting Information

**Table S1.** ΔH_MM-GBSA_ calculations and standard deviations averaged over all trajectory frames for all hit compounds during 100 ns.

| Ligand ID | 1<sup>st</sup> Replica<br>$\Delta H_{\text{MM-GBSA}}$<br>(kcal/mol) | 2<sup>nd</sup> Replica<br>$\Delta H_{\text{MM-GBSA}}$<br>(kcal/mol) | 3<sup>rd</sup> Replica<br>$\Delta H_{\text{MM-GBSA}}$<br>(kcal/mol) |
| --- | --- | --- | --- |
| G700-0411 |  |  | -44.06 ± 7.41 |
| P101-1799 | -43.63 ± 7.92 | -31.36 ± 8.77 | -39.20 ± 5.73 |
| S986-0170 |  | -54.05 ± 8.76 | -34.41 ± 6.47 |
| ZINC000096491549 | -49.85 ± 7.56 | -39.87 ± 6.02 |  |
| C336-0173 |  |  |  |
| ZINC000097784705 | -50.38 ± 4.28 |  | -55.27 ± 5.17 |
| ZINC000015062379 |  |  |  |
| ZINC000014882460 | -30.27 | -27.45 ± 7.46 |  |
| ZINC000091746401 | -36.08 ± 9.47 | -45.34 ± 7.37 | -23.31 ± 6.86 |
| ZINC000091965114 |  | -47.78 ± 4.62 | -45.59 ± 5.23 |

**Table S2.** ΔH_MM-GBSA_ calculations and standard deviations averaged over last 501 trajectory frames for all hit compounds during 100 ns.

| Ligand ID | 1<sup>st</sup> Replica<br>$\Delta H_{\text{MM-GBSA}}$<br>(kcal/mol) | 2<sup>nd</sup> Replica<br>$\Delta H_{\text{MM-GBSA}}$<br>(kcal/mol) | 3<sup>rd</sup> Replica<br>$\Delta H_{\text{MM-GBSA}}$<br>(kcal/mol) |
| --- | --- | --- | --- |
| G700-0411 | | | -43.33 ± 4.98 |
| P101-1799 | -40.91 ± 9.66 | -29.23 ± 7.69 | -40.12 ± 4.50 |
| S986-0170 | | -53.11 ± 8.49 | -33.34 ± 4.73 |
| ZINC000096491549 | -48.80 ± 7.15 | -40.46 ± 5.84 | |
| C336-0173 |  |  |  |
| ZINC000097784705 | -49.19 ± 3.41 | | -56.85 ± 4.87 |
| ZINC000015062379 |  |  |  |
| ZINC000014882460 | -27.24 ± 8.35 | -28.60 ± 8.44 | |
| ZINC000091746401 | -42.24 ± 5.09 | -46.83 ± 7.25 | -22.20 ± 7.48 |
| ZINC000091965114 | | -47.35 ± 5.51 | -45.24 ± 4.61 |

